# Exploiting toxin internalization receptors to enhance delivery of proteins to lysosomes for enzyme replacement therapy

**DOI:** 10.1101/2020.01.22.915298

**Authors:** Seiji N. Sugiman-Marangos, Greg L. Beilhartz, Xiaochu Zhao, Dongxia Zhou, Rong Hua, Peter K. Kim, James M. Rini, Berge A. Minassian, Roman A. Melnyk

## Abstract

Lysosomal storage diseases are a group of over 70 inherited genetic diseases caused by a defect or deficiency in a lysosomal protein. Enzyme replacement therapy, in which a functional copy of the defective enzyme is injected either systemically or directly into the brain of affected individuals, has proven to be an effective strategy for treating certain lysosomal storage diseases; however, the inefficient uptake of recombinant enzymes into cells and tissues *via* the low-affinity mannose-6-phosphate receptor prohibits broader utility of replacement therapy. Here, to improve the efficiency and efficacy of lysosomal enzyme uptake, we exploited the strategy used by diphtheria toxin to enter into the endo-lysosomal network of cells by creating a chimera between the receptor-binding fragment of diphtheria toxin and the lysosomal hydrolase TPP1. We show that the targeted TPP1 chimera binds with high affinity to target cells and is delivered into lysosomes with much greater efficiency than TPP1 alone. Further, we demonstrate efficient and durable uptake of the chimera *in vivo* following intracerebroventricular injection in mice lacking TPP1. Targeting the highly efficient diphtheria toxin internalization pathway represents a novel approach for improving the efficacy and utility of enzyme replacement therapy for treating lysosomal storage diseases.

## Introduction

Lysosomal storage diseases (LSDs) are a group of more than 70 inherited childhood diseases characterized by an accumulation of cellular metabolites due to deficiencies in a specific protein, typically a lysosomal hydrolase. Although each individual disease is considered rare, all together LSDs have a combined incidence of between 1/5,000 to 1/8,000 live births, and account for a significant proportion of neurodegeneration diagnoses in children.^1^ The particular age of onset for a given LSD varies depending on the affected protein and the percentage of enzymatic activity still present; however, in most cases, symptoms manifest early in life and progress insidiously, affecting multiple tissues and organs.^2^ In all but the mildest of cases, disease progression results in severe physical disability, possible intellectual disability, and a shortened life expectancy, with death occurring in late childhood or early adolescence.

As monogenic diseases, which are caused by functionally absent lysosomal enzymes, re-introducing a functional form of the defective enzyme back into lysosomes is a simple and effective strategy to treat LSDs. In practice, the current approach for enzyme replacement therapy (ERT) typically involves bi-weekly intravenous infusions of the functional form of the deficient protein responsible for diseases. ERT is currently approved for treatment of seven LSDs, and clinical trials are ongoing for five others.^3^ An essential requirement for efficient rescue and treatment of LSD is the ability to deliver curative doses of recombinant lysosomal enzymes into the lysosomal compartment of cells. ERT takes advantage of a specific N-glycan post-translational modification, mannose-6-phosphorylation (M6P), which results in the uptake of lysosomal enzymes from circulation by cells possessing the cation-independent M6P-receptor (CIMPR).^4^ While ERT is an effective treatment for some somatic symptoms, a combination of factors including: the abundance of M6PR in the liver; poor levels of expression in several key tissue types such as bone and skeletal muscle; incomplete and inconsistent M6P labeling of recombinant enzymes; and the relatively poor affinity of M6P for its cell-surface receptor (*viz.* K_d_ = 7 μM^5^) – all contribute to diminishing the overall effectiveness of therapies using CIMPR for cell entry.^3^

To improve the delivery and efficacy of therapeutic lysosomal enzymes, we drew inspiration from bacterial toxins, which as part of their mechanism hijack specific host cell-surface receptors that allow them to be efficiently trafficked into the endo-lysosomal pathway. While we and others have explored exploiting this pathway to deliver cargo into the cytosol before it reaches the lysosomes,^6, 7^ here we asked whether this same pathway could be exploited to enhance the delivery of lysosomal enzymes into lysosomes to improve ERT. Specifically, we sought to target the diphtheria toxin (DT) receptor owing to its high expression on neurons, the efficiency of its uptake and the lack of endogenous ligand that could present mechanism-based toxicity for other toxin-ligand combinations.

*Corynebacterium diphtheriae* secretes DT exotoxin, which is spread to distant organs by the circulatory system affecting the lungs, heart, liver, kidneys and the nervous system.^8^ It is estimated that 75% of individuals with acute disease also develop some form of peripheral or cranial neuropathy. This multi-organ targeting is a result of the ubiquitous nature of the DT receptor, Heparin-Binding EGF-like Growth Factor (HBEGF).

Diphtheria toxin is a three-domain protein that consists of an amino-terminal ADP-ribosyl transferase enzyme (A), a central translocation domain (T), and a carboxy-terminal receptor-binding moiety (R), which is responsible for binding cell-surface HBEGF with high affinity and triggering endocytosis into early endosomes (**Figure 1A**). Within endosomes, DT_T_ forms membrane-spanning pores that serve as conduits for DT_A_ to ‘escape’ from endosomes into the cytosol where it inactivates the host protein synthesis machinery. The remaining majority of toxin that does not escape endosomes continues to lysosomes where it is degraded.^9, 10^ We hypothesized that the receptor-binding domain, lacking any means to escape endosomes, would proceed with any attached cargo to lysosomes and thus serve as a means to deliver cargo specifically into lysosomes following high affinity binding to HBEGF.

**Figure 1.**
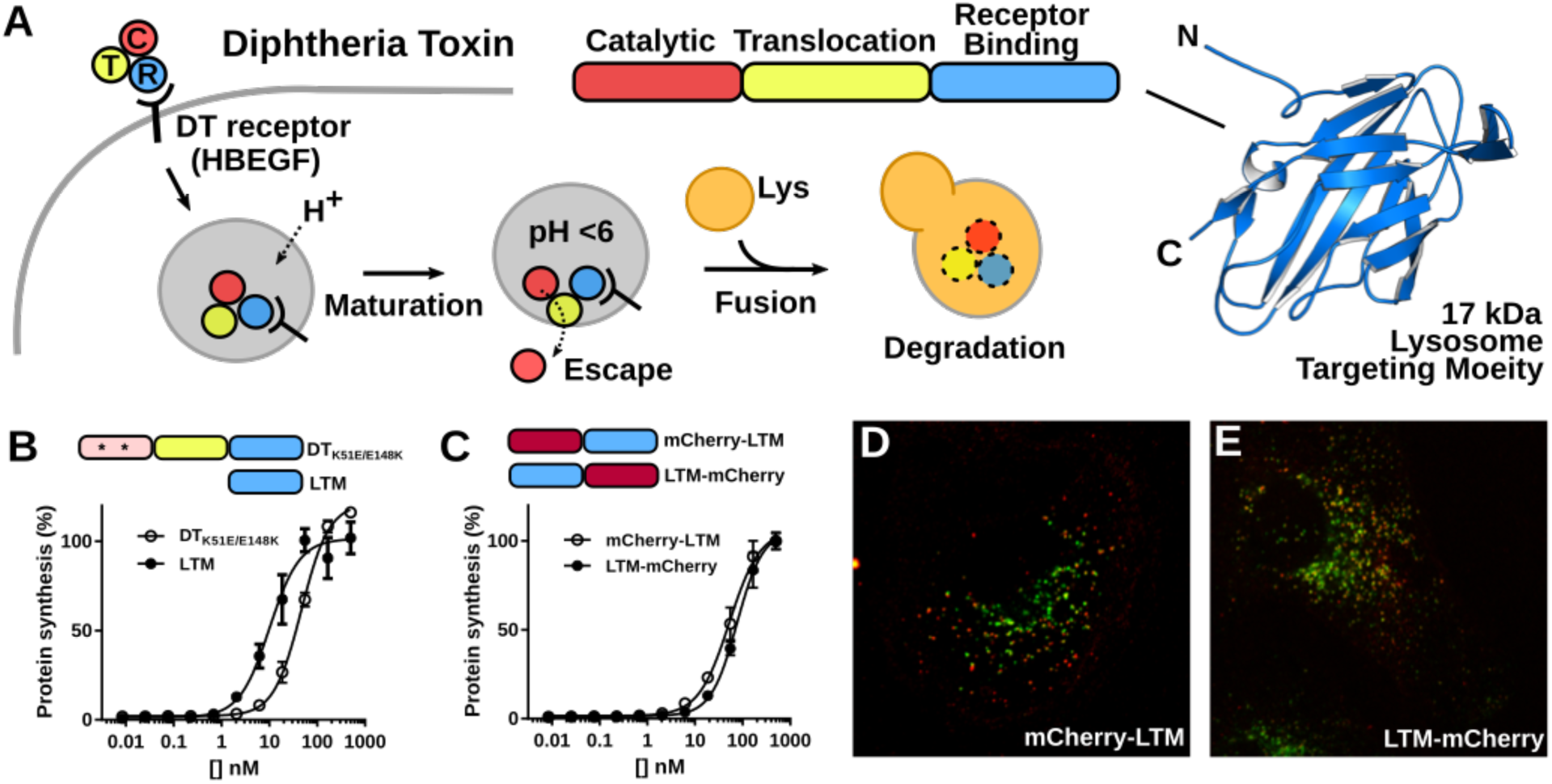
LTM fusion proteins bind HBEGF and traffic to the lysosome. (A) DT intoxication pathway (left), DT domain architecture (right). (B,C) DT_K51E/E148K_, LTM, mCherry-LTM and LTM-mCherry compete with wild-type diphtheria toxin for binding and inhibit its activity in a dose dependent manner with IC_50_’s of 46.9 nM, 10.1 nM, 52.7 nM and 76.1 nM, respectively (mean ± SD, n=3). (D,E) C-terminal and N-terminal fusions of LTM to mCherry were immuno-stained (red) and observed to co-localize with the lysosomal marker LAMP-1 (green).

In this study, we generated a series of chimeras containing the DT receptor-binding moiety, DT_R_, with the goal of demonstrating the feasibility of delivering therapeutic enzymes into lysosomes through a novel internalization pathway. We show that DT_R_ serves as a highly effective and versatile lysosomal targeting moiety (LTM) at either terminus of cargo, retaining its high affinity binding to HBEGF and trafficking into lysosomes both *in vitro* and *in vivo*. Based on the practical and fundamental advantages offered by DT_R_-mediated delivery of cargo into lysosome, over M6P-mediated mechanisms, we investigated the utility of DT_R_ as as a lysosomal delivery platform via fusions to human tripeptidyl peptidase-1 (TPP1) enzyme as an enhanced enzyme replacement approach for Batten disease.

## Results

### The DT receptor binding fragment is an autonomous Lysosome-Targeting Moiety (LTM)

To evaluate whether the diphtheria toxin receptor binding fragment could function autonomously to traffic cargo into lysosomes, we first asked whether the isolated 17-kDa DT_R_ fragment could be expressed independently from the DT holotoxin and retain its affinity for HBEGF. We cloned, expressed and purified the receptor-binding fragment and evaluated its ability to compete with full-length diphtheria toxin for the diphtheria toxin receptor, HBEGF. Prior to treating cells with a fixed dose of wild-type DT that completely inhibits protein synthesis, cells were incubated with a range of concentrations of DT_R_ or a full-length, non-toxic mutant of DT (DT_K51E_/_E148K_). DT_R_-mediated inhibition of wild-type DT-mediated toxicity was equivalent to nontoxic DT (**Figure 1B**), demonstrating that the receptor-binding fragment can be isolated from the holotoxin without affecting its ability to fold and bind cell-surface HBEGF. Next, we evaluated whether DT_R_ had a positional bias (*i.e.*, was able to bind HBEGF with a fusion partner when positioned at either terminus). To this end, we generated N- and C-terminal fusions of LTM to the model fluorescent protein mCherry (*i.e.,* mCherry-LTM and LTM-mCherry). To determine binding of each chimera to HBEGF, we quantified the ability of each chimera to compete with wild-type DT on cells in the intoxication assay. Both constructs competed with wild-type DT to the same extent as LTM alone and DT_K51E/E148K_ (**Figure 1C**), demonstrating that the LTM is versatile and autonomously folds in different contexts.

To evaluate intracellular trafficking, HeLa cells were treated with either LTM-mCherry or mCherry-LTM and then fixed and stained with an antibody against the lysosomal marker LAMP-1. In both cases, we observed significant uptake of the fusion protein and co-localization of mCherry and LAMP-1 staining (**Figures 1D and E**) indicating trafficking to the lysosomal compartments of the cells. Taken together, these results confirm that the LTM is capable of binding HBEGF independent of the holotoxin and trafficking associated cargo into cells and that the LTM can function in this manner at either terminus of a fusion construct.

### LTM-TPP1 retains TPP1 enzymatic activity and HBEGF binding

With no positional bias observed, we next screened LTM fusions to TPP1 to identify a design that maximizes expression, stability, activity and ultimately delivery. TPP1 is a 60-kDa lysosomal serine peptidase with a substrate preference for N-terminal tripeptides. Exposure to the low pH environment of the lysosome triggers auto-proteolytic activation, and the release of a 20-kDa pro-peptide which occludes its active site.^11^ From a design perspective, we favored an orientation in which the LTM was amino-terminal of TPP1, as auto-processing of TPP1 would result in the release of the upstream LTM-TPP1_propeptide_, liberating active, mature TPP1 enzyme in the lysosome (**Figure 2A**). Given the need for mammalian expression of lysosomal enzymes, we initially generated synthetic genetic fusions of the LTM to TPP1 in which we converted the codons from bacterially derived DT into the corresponding mammalian codons. HEK293F suspension cells stably expressing recombinant TPP1 (rTPP1) and TPP1 with an N-terminal LTM fusion (LTM-TPP1) were generated using the PiggyBac transposon system.^12^ A C-terminal construct (TPP1-LTM) was also produced; however, expression of this chimera was poor in comparison to rTPP1 and LTM-TPP1 (~0.4 mg/L cf. 10-15 mg/L).

**Figure 2.**
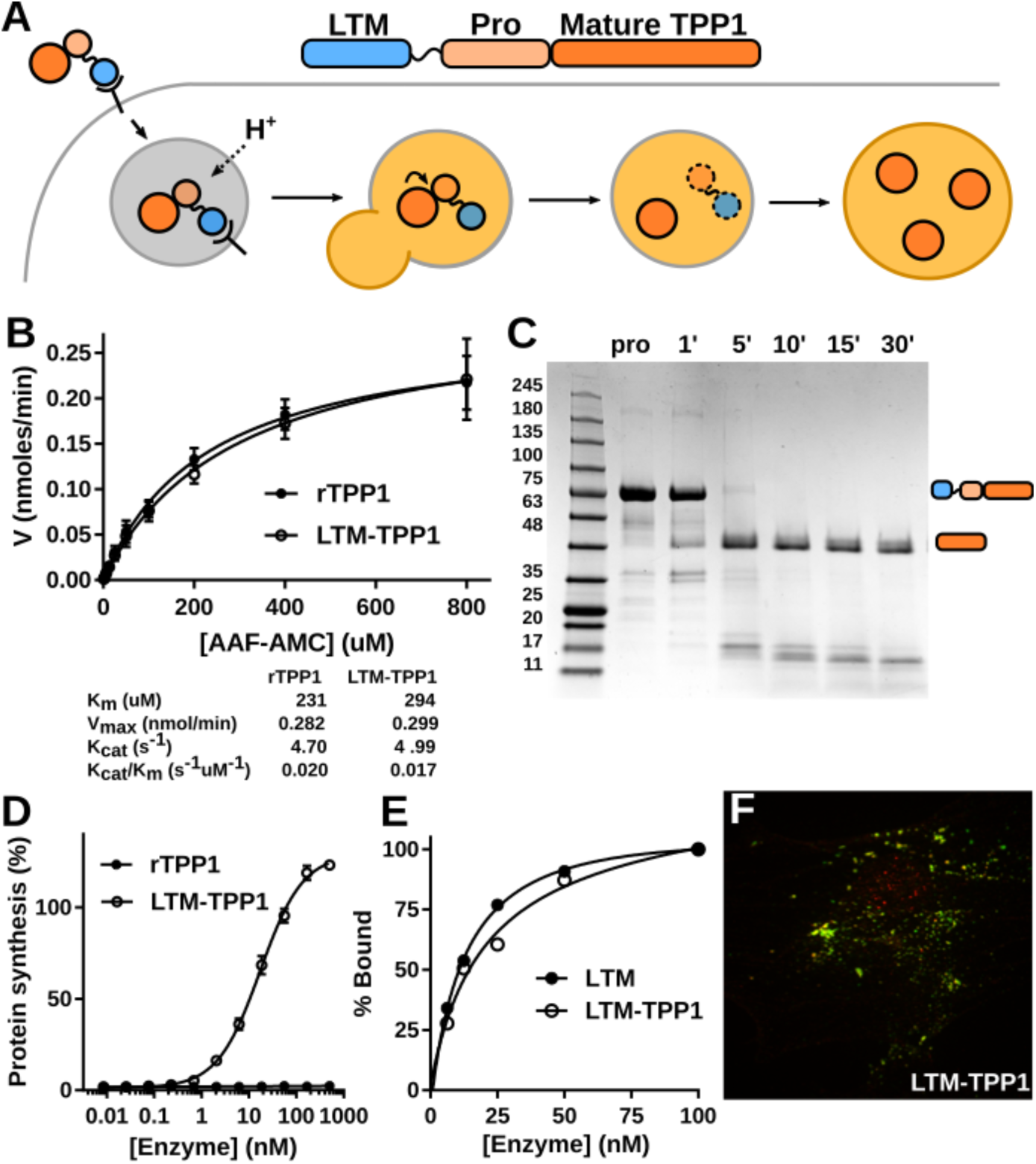
LTM and TPP1 fusion do not affect each other’s activity. (A) Design of LTM-TPP1 fusion protein and delivery schematic. (B) Enzyme kinetics of rTPP1 and LTM-TPP1 against the synthetic substrate AAF-AMC are indistinguishable. Michaelis-menten plots were generated by varying [AAF-AMC] at a constant concentration of 10 nM enzyme (mean ± SD, n=3). Plots and kinetic parameters were calculated with GraphPad Prism 7.04. (C) Maturation of TPP1 is unaffected by N-terminal fusion of LTM. (D) LTM-TPP1 inhibits wild-type DT activity in a dose dependent manner (IC_50_ of 17.2 nM) while rTPP1 has no effect on protein synthesis inhibition by DT (mean ± SD, n=3). (E) LTM and DTR-TPP1 bind HBEGF with apparent K_d_’s of 13.3 nM and 19.1 nM respectively. (F) LTM-TPP1 (green) co-localizes with LAMP-1 staining (red).

The activity of rTPP1 and LTM-TPP1 against the tripeptide substrate Ala-Ala-Phe-AMC were assessed to determine any effects of the LTM on TPP1 activity. The enzyme activities of rTPP1 and LTM-TPP1 were determined to be equivalent as evidenced through measurements of their catalytic efficiency (**Figure 2B**), demonstrating that there is no inference by LTM on the peptidase activity of TPP1. Maturation of LTM-TPP1 through auto-catalytic cleavage of the N-terminal pro-peptide was analyzed by SDS-PAGE (**Figure 2C**). Complete processing of the zymogen at pH 3.5 and 37°C occurred between 5 and 10 minutes, which is consistent with what has been observed for the native recombinant enzyme.^13^

The ability of LTM-TPP1 to compete with DT for binding to extracellular HBEGF was first assessed with the protein synthesis competition assay. Like LTM, mCherry-LTM and LTM-mCherry, LTM-TPP1 prevents protein synthesis inhibition by 10 pM DT with an IC_50_ of 17.2 nM (**Figure 2D**). As expected, rTPP1 alone was unable to inhibit DT-mediated entry and cytotoxicity. To further characterize this interaction, we measured the interaction between LTM and LTM-TPP and recombinant HBEGF using suface plasmon resonance binding analysis (**Figure 2E**). By SPR, the LTM and LTM-TPP1 were calculated to have apparent dissociation constants (K_d_) of 13.3 nM and 19.1 nM respectively, closely tracking the IC_50_ values obtained from the competition experiments (10.1 nM and 17.2 nM, respectively). Consistent with these results, LTM-TPP1 co-localizes with LAMP1 by immunofluorescence (**Figure 2F**).

### CRISPR-Cas9 mediated knockout of CLN2 in HeLa Kyoto cells

To study uptake of chimeric fusion proteins in cell culture, we generated a cell line deficient in TPP1 activity. A CRISPR RNA was designed targeting the signal peptide region of TPP1 in exon 2 of CLN2. Human HeLa Kyoto cells were reversed transfected with a Cas9 ribonucleoprotein complex and then seeded at low density into a 10 cm dish. Single cells were expanded to colonies which were picked and screened for TPP1 activity. A single clone deficient in TPP1 activity was isolated and expanded which was determined to have ~4% TPP1 activity relative to wild-type HeLa Kyoto cells plated at the same density (**Figure 3A**). The small residual activity observed is likely the result of another cellular enzyme processing the AAF-AMC substrate used in this assay as there is no apparent TPP1 protein being produced (**Figure 3B**). Sanger sequencing of the individual alleles confirmed complete disruption of the CLN2 gene (**Supplemental Figure 1**). In total, 3 unique mutations were identified within exon 2 of CLN2: a single base insertion resulting in a frameshift mutation; and two deletions of 24 and 33 bp respectively.

**Figure 3.**
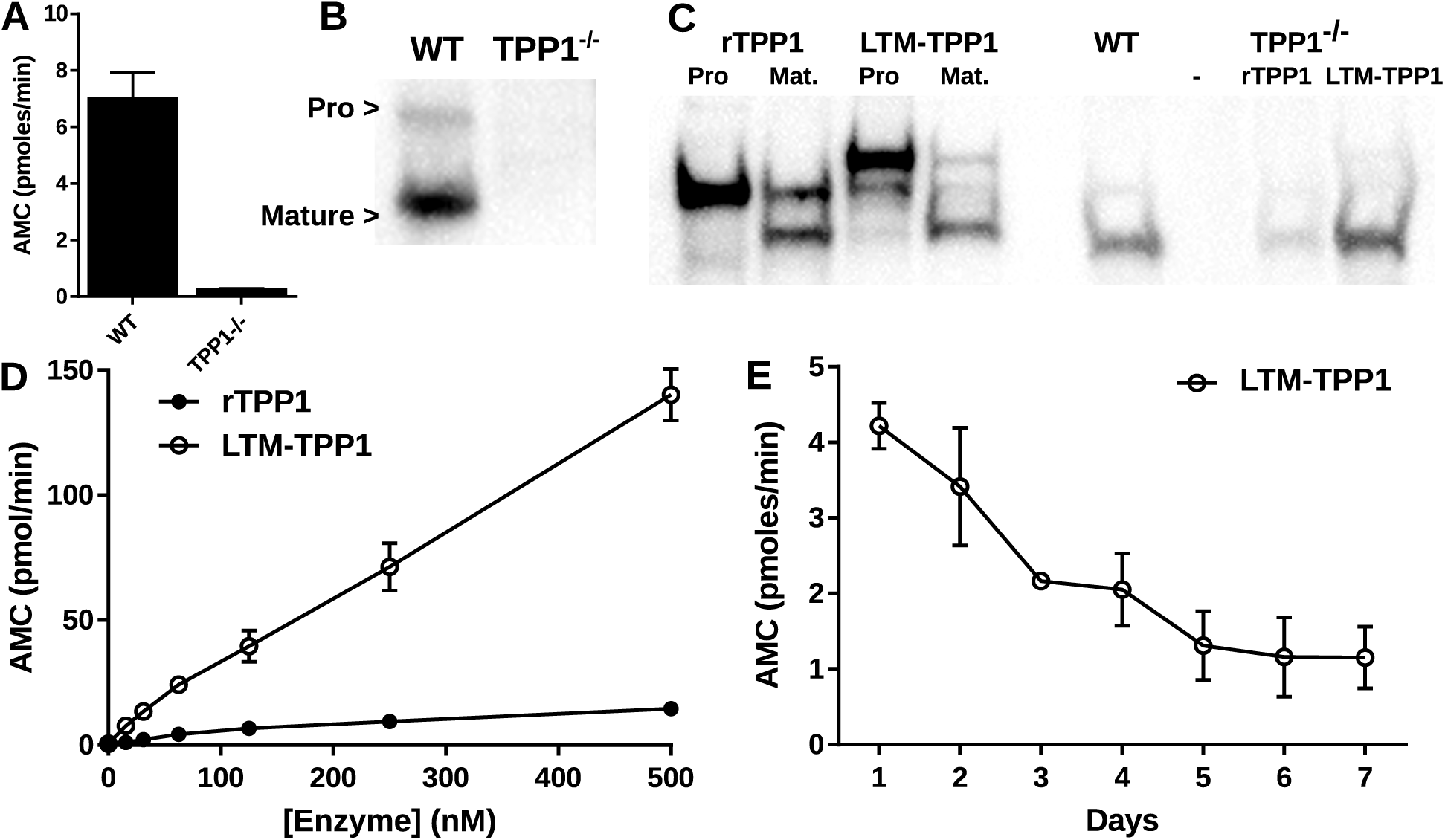
Intracellular uptake of LTM-TPP1 into CLN2 knockout cells. (A) CLN2 knockout cells exhibit ~4% TPP1 activity relative to wild-type HeLa Kyoto cells (mean ± SD, n=3). (B) Western blotting against TPP1 reveals no detectable protein in the knockout cells. (C)(*left*) In vitro maturation of pro-rTPP1 and LTM-TPP1 (16 ng) was analyzed by western blot. (*right*) TPP1 present in WT and TPP1^−/−^ cells, and TPP1^−/−^ cells treated with 100 nM rTPP1 and LTM-TPP1. (D) Uptake of rTPP1 and LTM-TPP1 into HeLa Kyoto TPP1^−/−^ cells was monitored by TPP1 activity (mean ± SD, n=4). (E) TPP1 activity present in HeLa Kyoto TPP1^−/−^ cells following a single treatment with 50 nM LTM-TPP1 (mean ± SD, n=3).

### LTM-TPP1 uptake into cells is more efficient than free rTPP1

Next, we compared the delivery and activation of rTPP1 and LTM-TPP1 into lysosomes by treating TPP1^−/−^ cells with a fixed concentration of enzyme (100 nM) and analyzing entry and processing by Western blot (**Figure 3C**). In both cases, the majority of enzyme was present in the mature form indicating successful delivery to the lysosome; however, the uptake of LTM-TPP1 greatly exceeded uptake of rTPP1. To quantify the difference in uptake and lysosomal delivery, cells were treated overnight with a range of doses of each enzyme, washed, lysed and assayed for TPP1 activity. Activity assays were performed without a pre-activation step, so signal represents protein that has been activated in the lysosome. For both constructs, we observed a dose-dependent increase in delivery of TPP1 to the lysosome (**Figure 3D**). Delivery of LTM-TPP1 was significantly enhanced compared to TPP1 alone at all doses, further demonstrating that uptake by HBEGF is more efficient than CIMPR alone. TPP1 activity in cells treated with LTM-TPP1 was consistently ~10x greater than that of cells treated with rTPP1, with the relative difference increasing at the highest concentrations tested. This may speak to differences in abundance, replenishment and/or recycling of HBEGF versus CIMPR. Uptake of LTM-TPP1 and rTPP1 into several other cell types yielded similar results (**Supplemental Figure 2**). To assess the lifetime of delivered enzyme, cells were treated with LTM-TPP1 (50 nM) and incubated overnight. Cells were washed and incubated with fresh media and TPP1 activity was assayed over the course of several days. Cells treated with LTM-TPP1 still retained appreciable TPP1 activity at 1-week post treatment (**Figure 4E**).

**Figure 4.**
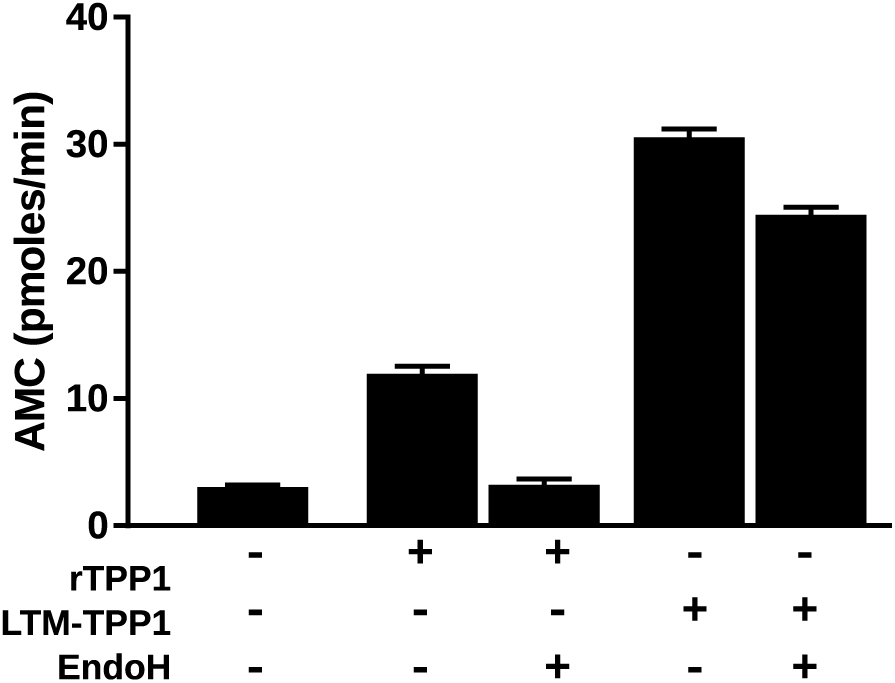
Uptake of rTPP1 and LTM-TPP1 into HeLa Kyoto TPP1^−/−^ cells following EndoH treatment to remove N-glycan labeling (mean ± SD, n=3).

While the DT competition experiment demonstrated that HBEGF is involved in the uptake of LTM-TPP1 but not rTPP1 (**Figure 2D**), it does not account for the contribution of CIMPR to uptake. Endoglycosidase H (EndoH) cleaves between the N-acetylglucosamine residues of N-linked glycans, leaving behind a single N-acetylglucosamine on the asparagine. Both rTPP1 and LTM-TPP1 were treated with EndoH to remove any M6P moieties and delivery into Hela TPP1^−/−^ was subsequently assessed. While rTPP1 uptake is completely abrogated by treatment with EndoH, LTM-TPP1 uptake is only partially decreased (**Figure 4**), indicating that while HBEGF mediated endocytosis is the principal means by which LTM-TPP1 is taken up into cells, uptake via CIMPR still occurs. The fact that CIMPR uptake is still possible in the LTM-TPP1 fusion means that the fusion is targeted to two receptors simultaneously, increasing its total uptake and potentially its biodistribution.

### Intracerebroventricular injection of LTM-TPP1 is taken up more efficiently into murine brain tissue than TPP1 alone

Uptake of LTM-TPP1 via the combination of HBEGF and CIMPR was shown to be 3-20x more efficient than CIMPR alone *in cellulo* (**Supplemental Figure 2**). To interrogate this effect *in vivo*, TPP1 deficient mice (TPP1^tm1pLob^ or TPP1^−/−^) were obtained as a kind gift from Dr. Peter Lobel at Rutgers University. Targeted disruption of the CLN2 gene was achieved by insertion of a neo cassette into intron 11 in combination with a point mutation (R446H), rendering these mice TPP1 null by both western blot and enzyme activity assay.^14^ Prior studies have demonstrated that direct administration of rTPP1 into the CSF via ICV or intrathecal (IT) injection results in amelioration of disease phenotype^15^ and even extension of lifespan in the disease mouse.^16^ To compare uptake of LTM-TPP1 and rTPP1 *in vivo*, the enzymes were injected into the left ventricle of 6-week old TPP1−/− mice. Mice were sacrificed 24 hours after injection and brain homogenates of wild-type littermates, untreated, and treated mice were assayed for TPP1 activity (**Figure 5A**). Assays were performed without pre-activation and therefore represent enzyme that has been taken up into cells, trafficked to the lysosome and processed to the mature form.

**Figure 5.**
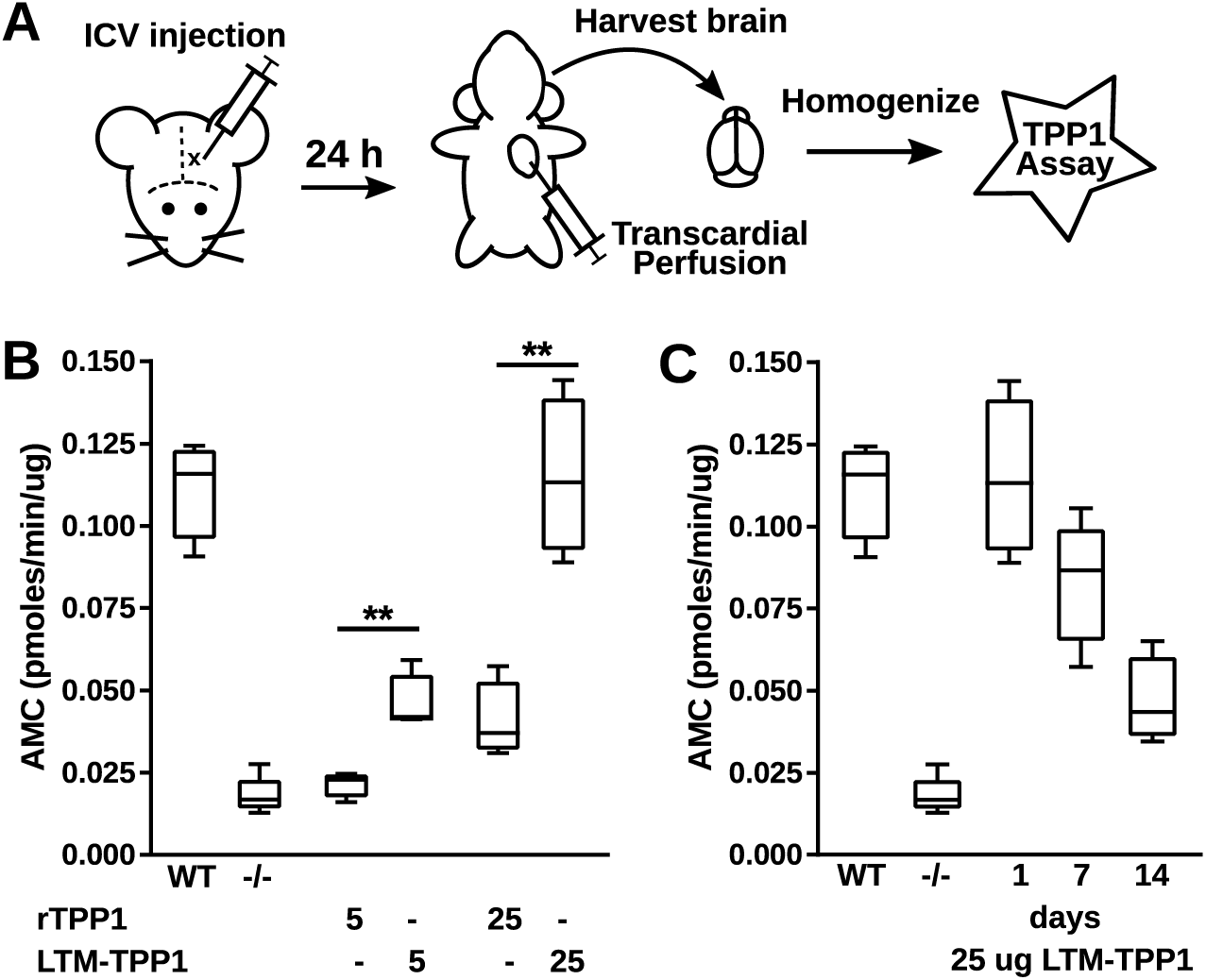
In vivo uptake of LTM-TPP1 following ICV injection. (A) Assay schematic. (B) TPP1 activity in brain homogenates of 6-week old mice injected with two doses (5ug, 25 ug) of either rTPP1 or LTM-TPP1 (5 ug–p=0.01, 25 ug–p=0.002). (C) TPP1 activity in brain homogenates following a single 25 ug dose of LTM-TPP1, 1, 7 and 14-days post-injection. Data is presented as box-and-whisker plots with whiskers representing min and max values from n≥4 mice per group. Statistical significance was calculated using paired t-tests with GraphPad Prism 7.04.

While both enzymes resulted in a dose dependent increase in TPP1 activity, low (5 ug) and high (25 ug) doses of rTPP1 resulted in only modest increases of activity, representing ~6% and ~26% of wild-type levels of activity, respectively (**Figure 5B**). At the same doses LTM-TPP1 restored ~31% and ~103% of wild-type activity. To assess the lifetime of enzyme in the brain, mice were injected ICV with 25 μg of LTM-TPP1 and sacrificed either 1- or 2-weeks post-injection. Remarkably, at 1-week post-injection, ~68% of TPP1 activity was retained (compared to 1-day post-injection), and after 2-weeks, activity was reduced to ~31% (**Figure 5C**).

## Discussion

ERT is a lifesaving therapy that is principal method of treatment in non-neurological LSDs. Uptake of M6P labelled enzymes by CIMPR is relatively ineffective due to low receptor affinity,^5^ heterogeneous expression of the receptor, and incomplete labeling of recombinantly produced enzyme.^17^ Despite its inefficiencies and high cost (~$200,000 USD/patient/year)^18^ it remains the standard of care for several LSDs, as alternative treatment modalities (substrate reduction therapy, gene therapy, hematopoetic stem cell transplantation) are not effective, not as well developed, or inherently riskier.^19–23^ Improving the efficiency and distribution of recombinant enzyme uptake may help address some of the current shortcomings in traditional ERT.

Several strategies have been employed to increase the extent of M6P labeling on recombinantly produced lysosomal enzymes: engineering mammalian and yeast cell lines to produce more specific/uniform N-glycan modifications,^17, 24, 25^ chemical or enzymatic modification of N-glycans post-translationally,^26^ and covalent coupling of M6P^27^. Some of these strategies have made incremental improvements to uptake but do nothing to address the relatively poor affinity of M6P for its receptor which is in the low uM range.^5^ M6P independent uptake of a lysosomal hydrolase by CIMPR has been previously demonstrated with beta-glucuronidase (GUS)^28^ and acid α-glucosidase (GAA).^29^ In the latter work, a peptide tag (GILT) targeting insulin-like growth factor II receptor (IGF2R) was fused to recombinant GAA. CIMPR is a ~300 kDa, 15-domain membrane protein with 3 M6P binding domains and 1 IGF2R domain. By targeting the EGF2R domain with a high-affinity (low nM) peptide rather than the low affinity M6P binding domain, the authors were able to demonstrate a >20-fold increase in uptake of their chimera in cell culture, and a ~5-fold increase in the ability to clear built-up muscle glycogen in GAA deficient mice.

In this study, we have demonstrated efficient uptake and lysosomal trafficking of a model lysosomal enzyme, TPP1, via a CIMPR independent route, using the receptor binding domain of a bacterial toxin. HBEGF is a mitogenic glycoprotein, a member of the EGF family of growth factors, and is the unique receptor for DT. Speaking to its global presence, it plays roles in cardiac development, wound healing, muscle contraction and neurogenesis, however, it does not act as a receptor in any of these physiological processes.^30^ Intracellular intoxication by DT is the only known process in which HBEGF acts as a receptor, making it an excellent candidate receptor for ERT as there is no natural ligand with which to compete. Upon binding, DT is internalized via clathrin-mediated endocytosis and then trafficked towards lysosomes for degradation.^31, 32^ Acidification of endosomal vesicles by vacuolar ATPases promotes the insertion of DT_T_ into the endosomal membrane and the subsequent translocation of the catalytic DT_A_ domain into the cytosol. In the absence of an escape mechanism, the majority of internalized LTM should be trafficked to the lysosome as we have demonstrated with our chimera (**Figure 3d, 4a,b,d**). Uptake of LTM-TPP1 in vitro is robust relative to rTPP1 (**Figure 4D, Supplemental Figure 2**) and TPP1 activity is sustained in the lysosome for a significant length of time (**Figure 4d**). We have also demonstrated that the increase in uptake efficiency that we observed in cell culture persists *in vivo*. TPP1 activity in the brains of CLN2 null mice was significantly greater in animals treated with ICV injected LTM-TPP1 as compared to this treated with TPP1 at two different doses (**Figure 5B**), and remarkably, this activity persists with an apparent half-life of ~8 days (**Figure 5C**).

While the ability of this construct to affect disease progression has yet to be determined, recent positive clinical trial results^33^ and the subsequent approval of recombinant TPP1 (cerliponase alfa) for treatment of Neuronal Ceroid Lipofuscinosis 2 (NCL2) provide support for this approach. In this trial, 300 mg of recombinant TPP1 was administered by bi-weekly ICV injection to 24 affected children and was able to stabilize their disease progression. While this dose is of the same order of magnitude as other approved ERTs (<1 to 40 mg/kg),^34, 35^ it represents a substantial dose especially considering that it is isolated to a single organ. Improving the efficiency of uptake by targeting an additional receptor as we have demonstrated and therefore decrease the dose necessary to improve symptoms would decrease costs as well as potential dose related side effects. Moreover, intravenous injection of recombinant TPP1 to treat somatic symptoms of the disease will encounter the same issues as other ERTs with regards to limitations imposed by M6P mediated uptake, and therefore strategies to improve uptake efficiency and biodistribution remain of great importance.

## Materials and Methods

### Protein expression and purification

DT_K51E/E148K_, LTM, LTM-mCherry, mCherry-LTM, and HBEGF constructs were cloned using the In-Fusion HD cloning kit (Clontech) into the Champion pET-SUMO expression system (Invitrogen). Recombinant proteins were expressed as 6His-SUMO fusion proteins in *Escherichia coli* (*E. coli*) BL21(DE3)pLysS cells. Cultures were grown at 37°C until an OD_600_ of 0.5, induced with 1 mM IPTG for 4 hours at 25°C. Cell pellets harvested by centrifugation were re-suspended in lysis buffer (20 mM Tris pH 8.0, 160 mM NaCl, 10 mM imidazole, lysozyme, benzonase and protease inhibitor cocktail) and lysed by 3 passages through an EmulsiFlex C3 microfluidizer (Avestin). Following clarification by centrifugation at 18,000xg for 20 minutes and syringe-filtration (0.2 um), soluble lysate was loaded over a 5 mL His-trap FF column (GE-Healthcare) using an AKTA FPLC. Bound protein was washed and eluted over an imidazole gradient (20-150 mM). Fractions were assessed for purity by SDS-PAGE, pooled, concentrated and frozen on dry-ice in 25% glycerol for storage at −80°C.

TPP1 cDNA was obtained from the SPARC biocentre (The Hospital for Sick Children) and cloned into the *piggyBac* plasmid pB-T-PAF (Jim Rini, University of Toronto) using NotI and AscI restriction sites to generate two expression constructs (pB-T-PAF-ProteinA-TEV-LTM-TPP1 and pB-T-PAF-ProteinA-TEV-TPP1). Stably transformed expression cell lines (HEK293F) were then generated using the *piggyBac* transposon system as described.^12^ Protein expression was induced with doxycycline and secreted fusion protein was separated from expression media using IgG Sepharose 6 Fast Flow resin (GE-Healthcare) in a 10 mL Poly-Prep chromatography column (Bio-Rad). Resin was washed with 50 column volumes of wash buffer (10 mM Tris pH 7.5, 150 mM NaCl) and then incubated overnight at 4°C with TEV protease to release the recombinant enzyme from the ProteinA tag. Purified protein was then concentrated and frozen on dry-ice in 50% glycerol for storage at −80°C.

### Competition assay

Cellular intoxication by DT was measured using a nanoluciferase reporter strain of Vero cells (Vero NlucP) as described previously.^7^ Briefly, Vero NlucP cells were treated with a fixed dose of DT at EC99 (10 pM) and a serial dilution of LTM, LTM-mCherry, mCherry-LTM, DT_K51E/E148K_, LTM-TPP1 or rTPP1 and incubated overnight (17 h) at 37°C. Cell media was then replaced with a 1:1 mixture of fresh media and Nano-Glo Luciferase reagent (Promega) and luminescence was measured using a SpectroMax M5e (Molecular Devices). Results were analyzed with GraphPad Prism 7.04.

### Surface Plasmon Resonance

SPR analysis was performed on a Biacore X100 system (GE Healthcare) using a CM5 sensor chip. Recombinant HBEGF was immobilized to the chip using standard amine-coupling at a concentration of 25 ug/mL in 10 mM sodium acetate pH 6.0 with a final response of 1000-2500 resonance units (RU). LTM and LTM-TPP1 were diluted in running buffer (200 mM NaCl, 0.02% Tween 20, 20 mM Tris pH 7.5) at concentrations of 6.25 nM to 100 nM and injected in the multi-cycle analysis mode with a contact time of 180 s and a dissociation time of 600 s. The chip was regenerated between cycles with 10 mM glycine pH 1.8. Experiments were performed in duplicate using two different chips. Binding data was analyzed with Biacore X100 Evaluation Software version 2.0.2, with apparent dissociation constants calculated using the 1:1 steady-state affinity model.

### Microscopy

HeLa cells were incubated with LTM-mCherry (0.5 uM), mCherry-LTM (0.5 uM) or LTM-TPP1 (2 uM) for 2 hours. Cells were washed with ice cold PBS, fixed with 4% paraformaldehyde, and permeabilized with 0.5% Triton X-100. mCherry constructs were visualized with a rabbit polyclonal antibody against mCherry (Abcam ab16745) and anti-rabbit Alexa Flour 568 (Thermo Fisher Scientific). LAMP1 was stained with a mouse primary antibody (DSHB 1D4B) and anti-mouse Alexa Fluor 488 (Thermo Fisher Scientific).

CLN2−/− fibroblast 19494 were incubated with LTM-TPP1 (2 uM) for 2 hours. Cells were washed with ice cold PBS, fixed with 4% paraformaldehyde, and permeabilized with 0.5% Triton X-100. LTM-TPP1 was visualized with a mouse monoclonal against TPP1 (Abcam ab54685) and anti-mouse Alexa Fluor 488 (Thermo Fisher Scientific). LAMP1 was stained with rabbit anti-LAMP1 and anti-rabbit Alexa Fluor 568 (Thermo Fisher Scientific).

### In vitro TPP1 activity assay

TPP1 protease activity was measured using the synthetic substrate Ala-Ala-Phe 7-amido-4-methlycoumarin (AAF-AMC) using a protocol adapted from Vines and Warburton.^36^ Briefly, enzyme was pre-activated in 25 uL activation buffer (50 mM NaOAc pH 3.5, 100 mM NaCl) for 1 hour at 37°C. Assay buffer (50 mM NaOAc pH 5.0, 100 mM NaCl) and substrate (200 uM AAF-AMC) were then added to a final volume of 100 uL. Fluorescence (380 nm excitation/460 nm emission) arising from the release of AMC was monitored in real-time using a SpectroMax M5e (Molecular Devices). TPP1 activity *in cellulo* was measured similarly, without the activation step. Cells in a 96-well plate were incubated with 25 uL of 0.5% Triton X-100 in PBS, which was then transferred to a black 96-well plate containing 75 uL of assay buffer with substrate in each well.

### Generation of CLN2^−/−^ cell line by CRISPR-Cas9

crRNA targeting the signal peptide sequence in exon 2 of CLN2 was designed using the Integrated DNA Technologies (www.idtdna.com) design tool. The gRNA:Cas9 ribonucleoprotein complex was assembled according to the manufacturer’s protocol (Integrated DNA technologies), and reverse transfected using Lipofectamine® RNAiMAX (Thermo Fisher Scientific) into HeLa Kyoto cells (40,000 cells in a 96-well plate). Following 48 hours incubation, 5,000 cells were seeded into a 10 cm dish. Clonal colonies were picked after 14 days and transferred to a 96-well plate. Clones were screened for successful CLN2-knockout by assaying TPP1 activity and confirmed by Sanger sequencing and Western blot against TPP1 antibody (Abcam ab54385).

### Auto-activation

The pro-form of TPP1 was matured *in vitro* to the active form in 50 mM NaOAc pH 3.5 and 100 mM NaCl for 1 to 30 minutes at 37°C. The auto-activation reaction was halted by the addition of 2x Laemmli SDS sample buffer containing 10% 2-mercaptoethanol and boiled for 5 minutes. Pro and mature TPP1 were separated by SDS-PAGE and imaged on a ChemiDoc™gel imaging system (Bio-rad)

### Western blot

Proteins or cellular lysate were separated by 4-20% gradient SDS-PAGE before being transferred to a nitrocellulose membrane using the iBlot™ (Invitrogen) dry transfer system. Membranes were then blocked for 1 hour with a 5% milk-TBS solution and incubated overnight at room temperature with a 1:1,00 dilution of mouse monoclonal antibody against TPP1 (Abcam ab54685) in 5% milk-TBS. Membranes were washed 3×5 min with 0.1% Tween® 20 (Sigma-Aldrich) in TBS before a 1 hour incubation with a 1:5,000 dilution of sheep anti-mouse IgG horseradish perodixase secondary antibody (GE Healthcare) in 5% milk-TBS. Chemiluminescent signal was developed with Clarity™ Western ECL substrate (Bio-rad) and visualized on a ChemiDoc™ gel imaging system (Bio-rad).

### Treatment with Endo H

rTTP1 and LTM-TPP1 were treated with Endoglycosidase H (New England Biolabs) to remove N-glycan modifications. Enzymes were incubated at 1 mg/mL with 2,500 units of Endo H for 48 hours at room temperature in 20 mM Tris pH 8.0, 150 mM NaCl in a total reaction volume of 20 uL. Cleavage of N-glycans was assessed by SDS-PAGE and concentrations were normalized to native enzyme specific activities.

### Animals

Cryo-preserved TPP1^+/−^ embryos were obtained from Dr. Peter Lobel at Rutgers University and re-derived in a C57/BL6 background at The Centre for Phenogenomics in Toronto. Animal maintenance and all procedures were approved by The Centre for Phenogenomics Animal Care Committee and are in compliance with the CCAC Guidelines and the OMAFRA Animals for Research Act.

### Intracerebroventricular injection

TPP1^−/−^ mice (60 days old) were anesthetized with isoflurane (inhaled) and injected subcutaneously with sterile saline (1 mL) and Meloxicam (2 mg/kg). Mice were secured to a stereotactic system, a small area of the head was shaved and single incision was made to expose the skull. A high-speed burr was used to drill a hole at stereotaxic coordinates: A/P -1.0 mm, M/L -0.3 mm, D/V -3.0 mm relative to the bregma, and a 33G needle attached to a 10 uL Hamilton syringe was used to perform the ICV injection into the left ventricle. Animals received either 1 uL or 5 uL of enzyme (5 ug/uL), injected at a constant rate. Isoflurane anesthetized animals were sacrificed by transcardial perfusion with PBS. Brains were harvested and frozen immediately, then thawed and homogenized in lysis buffer (500 mM NaCl, 0.5% Triton X-100, 0.1% SDS, 50 mM Tris pH 8.0) using 5 mM stainless steel beads in a TissueLyser II (Qiagen). In vitro TPP1 assay was performed as described, minus the activation step.

---

**Supplemental Figure 1.**
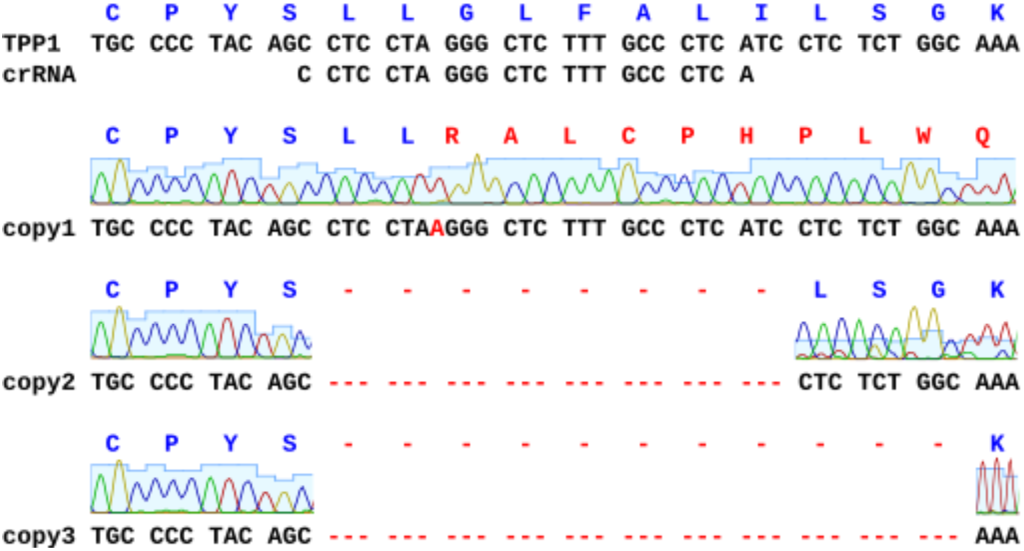
CRISPR-Cas9 knockout of CLN2.Sanger sequencing of genomic DNA from CLN2 knockout cells reveal 3 distinct mutations: a single-base insertion and 2 deletions.

**Supplemental Figure 2.**
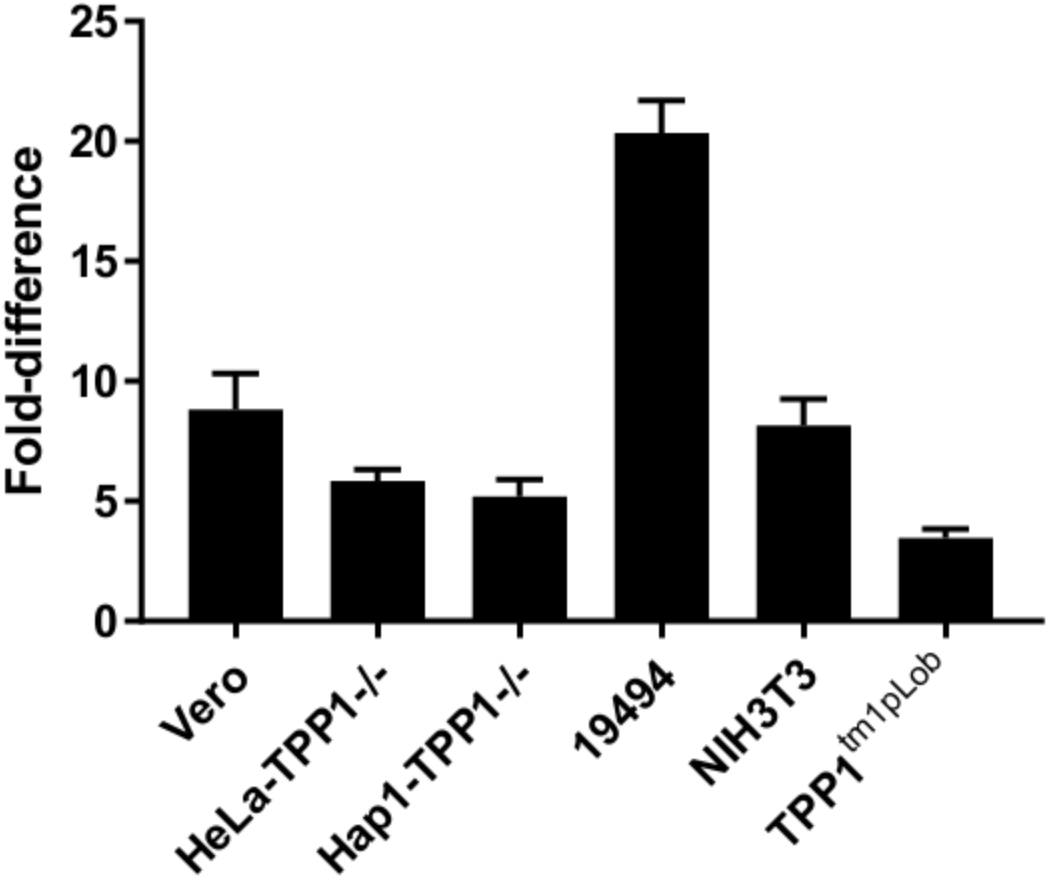
Relative uptake of rTPP1 vs. LTM-TPP1. Various cell types (Vero (African green monkey kidney epithelial), HeLa Kyoto TPP1−/− (human cervical cancer), Hap1 TPP1−/− (haploid human cells derived from KBM-7), 19494 (NCL2 patient fibroblasts), NIH3T3 (mouse fibroblasts), TPP1tm1pLob−/− (primary lung fibroblasts from TPP1 ko mouse)) were assayed for TPP1 activity following treatment with either rTPP1 or LTM-TPP1 (100 nM). Enzyme activity from cells treated with LTM-TPP1 was divided by activity of cells treated with rTPP1 to determine the relative difference in uptake efficiency for each cell type (mean ± SD, n=3).

